# Lactate-derived HDAC6 Lactylation as a new target for neuronal protection in cerebral ischemic reperfusion injury

**DOI:** 10.1101/2022.11.27.518054

**Authors:** Jun Cao, Zexuan Hong, Fanning Zeng, Zaisheng Qin, Miao Lin, Hao Wang, Daming Zuo, Tao Tao

## Abstract

Lactylation is a newly found lactate-derived post-translational modification, which is firstly reported on histone protein lysine residues to regulate gene expression. Here, we provided evidence that higher serum lactate level was possibly associated with better prognosis in ischemic stroke patients. Meanwhile, inhibition of endogenous lactate increased the infarct volume and decreased protein pan-lactylation in the affected brain tissue in cerebral ischemia mice. We further demonstrated that downregulating lactylation level of Hdac6 (Histone Deacetylase 6) in neurons exacerbated ischemic neuronal injury. The underlying mechanisms involved Hdac6 lactylation disturbing calcium homeostasis through binding BiP (Binding immunoglobulin protein), an endoplasmic reticulum-related protein. Collectively, our findings identified a new effect of lactate for neuroprotection through Hdac6 lactylation during cerebral ischemic reperfusion injury which introduced a potential therapeutic approach against ischemic stroke.

## Introduction

Lactate, an anaerobic glycolysis marker and supplemental energy fuel, now have been considered to play versatile roles in physiological and pathological conditions through regulating integrated physiological responses, cerebral blood flow, and neuronal excitability in the brain (Magistretti & Allaman, 2018). In cerebral ischemia, lactate increasing is the hallmark of energetic metabolism dysfunction, caused by the total or partial decrease in the oxygen and glucose supply after the sudden blood flow interruption (Bouzat & Oddo, 2014). A recent human study found that the level of cerebral lactate correlated with ultimate infarct volume in hypoperfused tissue through magnetic resonance spectroscopy imaging (MRSI) and better recovery was observed in the area with lower lactate level (Li *et al*, 2020). However, early investigations suggested that exogenous supplement lactate could produce neuroprotection in transient middle cerebral artery occlusion model (Berthet *et al*, 2009) and could increase energy availability in glutamate induced excitotoxicity of cultured cortical neurons (Jourdain *et al*, 2016). These seemingly paradoxical results raise the question of “To Lac or not to Lac” in cerebral ischemia. In 2019, the newly identified histone lactylation endowed the lactate a non-metabolic role, which may regulate genes expression through epigenetics. Lactate as a classical energetic molecule tangled in cerebral ischemia research for several decades.

Lactylation, a new post-translational medication (PTM) firstly found in histone lysine residues, ignites the thinking on lactate-induced PTM in biochemistry and widens the avenue of lactate research in physiological and pathological conditions (Zhang et al, 2019a). Lately, histone lactylation has been found as a new path to metabolic reprogramming and epigenetic modification in tumor (Sun et al, 2022). Moreover, stress-associated neural excitation induced histone H1 lactylation, which positively correlated with decreased social behavior (Hagihara et al, 2021). Increased histone lactylation has also been found in Alzheimer’s disease and the H4 lactylation aggravated the microglia dysfunction through stimulating glycolytic genes transcription (Pan et al, 2022). Additionally, histone lactylation also exhibited its protection in post-myocardial infarction by promoting reparative genes transcription in monocytes. Besides histone lactylation, a sepsis study reported that HMGB1 lactylation increasing its exocytosis from macrophage, which provided a new way to combat sepsis in lactate-related pathway (Yang et al, 2022). The latest report further confirmed that lactylation was widespread in the human proteome and probably played an important regulatory role (Wan et al, 2022). These findings provoke us to revisit the role of lactate and anticipate the effect of lactylation in cerebral ischemia.

Therefore, in current study, we examined the serum lactate level of cerebral ischemia patients and performed lactylation proteomic profiles in mice cerebral ischemic reperfusion model. We found that increased Hdac6 lactylation, a member of the Hdacs class IIb family, could reduce neuronal injury in ischemic reperfusion injury conditions via biding BiP, a calcium-related protein localized to endoplasmic reticulum.

## Result

### Endogenous lactate exerts neuroprotective effects by enhancing protein lactylation in cerebral ischemic reperfusion injury

We analyzed the clinical data of 30 patients with ischemic stroke retrospectively. Interestingly, patients with better neurological prognosis (modified Rankin Scale (mRS) ≤ 2) had significantly higher serum lactate than that with worse prognosis (mRS ≥ 3) (Fig1 A). In addition, serum lactate of all patients was higher than that of healthy volunteers (Fig1 B). Corresponding to patients, in tMCAO mice, downregulating endogenous lactate through oxamate intraperitoneal injection, significantly increased infarction volume compared to saline treatment group (Fig1 C). As indicated by the Immunofluorescence staining, the dendrites with lower endogenous lactate exhibited a more incoherent and disarranged structure in the ischemic penumbra neurons (Fig1 D). Western blot analysis in the affected brain hemisphere showed that Pan-lysine lactylation (Pan Kla) level was higher in tMCAO group compared to the sham group, or oxamate-treatment group in tMCAO (Fig1 E). Also, the immunofluorescence of affected brain slices exhibited the similar tendency of Pan Kla among three groups mentioned above (Fig1 F). To testify the endogenous effect of lactate on neuron in vitro, oxygen and glucose deprivation followed by reoxygenation (OGD/R) was conducted for primary cultured neuron with or without oxamate treatment. Under OGD/R, immunofluorescence staining reflected a more disorganized neuronal morphology in oxamate treated group compared with control group (Fig1 G). Meanwhile, TUNEL assay showed lower endogenous lactate accompanied with an increase in neuron death under OGD/R (Fig1 G).

**Fig 1.**
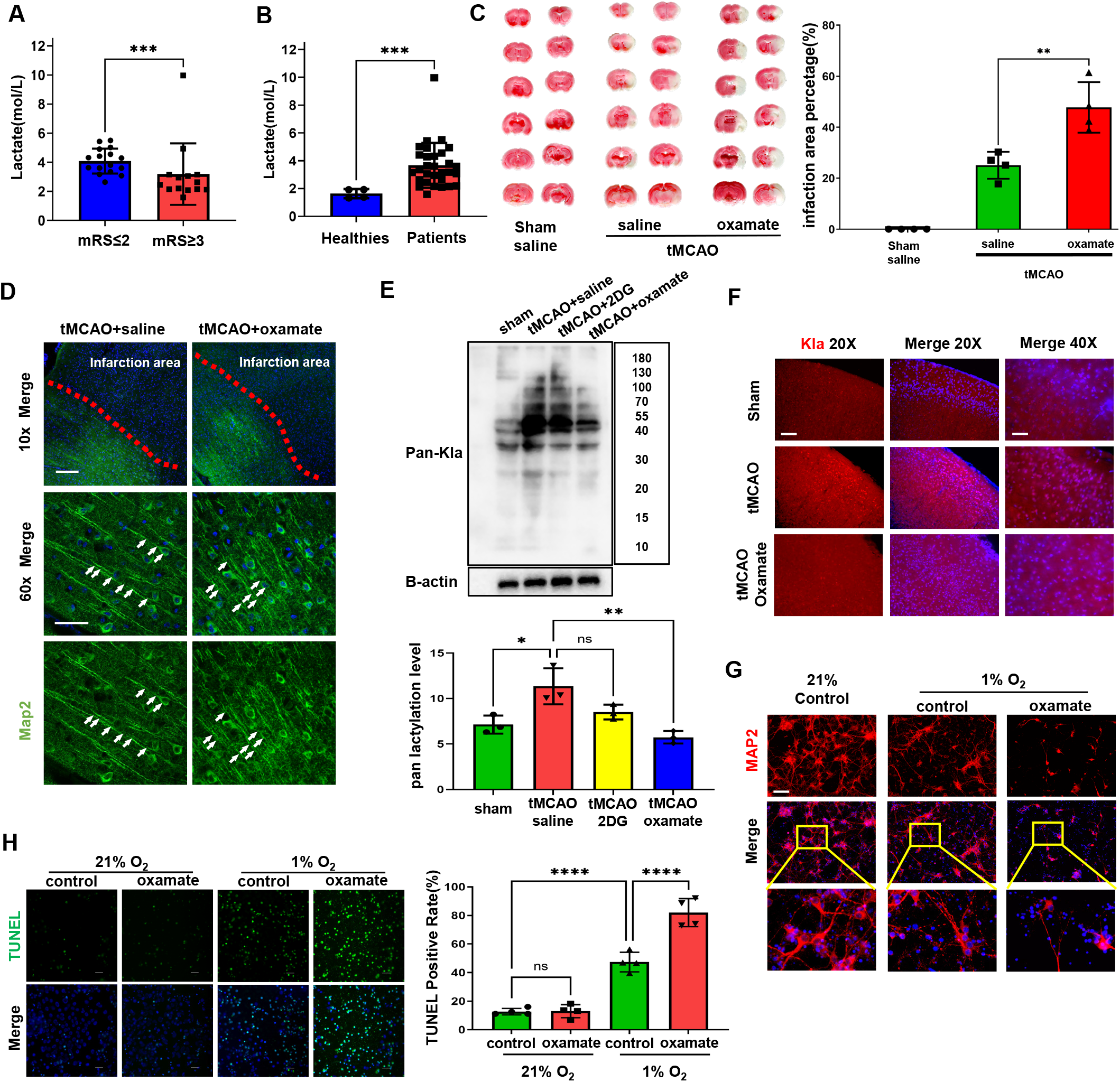
Lactate improved neuron functional outcomes and promoted protein lactalytion level in ischemic stroke. A. The serum lactate level in patients with better neurological prognosis (mRS≤ 2, n=16) or worse prognosis (mRS≥3, n=14); B. The lactate of serum from cerebral ischemia stroke patients (n=30) and healthy volunteers (n=4); C. TTC staining of the brain slices and the percentage of the infarct volume. (n=4); D. Immunostaining of Map2 in brain slices area. Arrows indicated the morphology of the neuron dendrites. Scale bar for 10X: 200μm; Scale bar for 60X : 50μm; E. Immunoblotting showed protein expression of Pan-Kla in vivo; (n=3) F. Brain slices immunofluorescence staining of Pan-Kla in ischemic penumbra. Scale bar for 20X: 100μm; Scale bar for 40X: 50μm; G. Immunostaining of Map2 in primary culture neurons with or without oxamate after OGD/R. Scale bar: 100μm; H. TUNEL staining of primary culture neurons with or without oxamate after OGD/R. (n=4) Scale bar: 50μm; Data in A,B are presented as median with interquartile range (Mann Whitney test); Data in C, E, H are presented as mean ± SD (Tukey’s multiple comparisons test); **\*** P<0.05, **\*\*** P<0.01, **\*\*\*\*** P<0.0001, ns P>0.05.

In all, these results indicated that endogenous lactate was possibly related to the clinical prognosis in stroke patients. The lactate-derived lactylation may involve in the neuroprotection by lactate in vivo and vitro.

### Lactylation of Hdac6 is relative to the neuronal injury in cerebral ischemic reperfusion injury

To explore the relationship between protein lactylation and the pathological mechanism of ischemic stroke, we analyzed enriched lactylated peptides of brain tissue in two groups (sham and tMCAO) using Liquid Chromatography Triple Quadrupole Mass Spectrometry (LC-MS/MS) search tools for resulting MaxQuant data. The heatmap in Figure EV1 A showed no significant difference in lactylation among the biological replicates within each group respectively. Figure EV1B showed that lactylation was mainly concentrated on lysine (K) residues of the proteins. LC-MS/MS analysis identified 204 lactylation proteins (Fig EV1C) with 160 upregulated and 49 downregulated lactylated sites. The GO network showed the outlook of functional connections and clustering analyses on the LC-MS/MS identified proteins (Fig 2A left). Since there are internal connections between each functional cluster to perform specific biological functions, we performed secondary clustering on all functional clusters to find out the functional subsets that were closely related to cerebral ischemia-reperfusion injury. The top1 subset of which mainly highlighted axon-synaptic-dendrites functions (Fig 2A right). The resting seven subsets were shown in Figure EV2A. Then the protein-protein interaction (PPI) of the top1 subset above was analyzed (Fig2 B), the top5 hub proteins from which were Hdac6, Mapt, Snca, Alb, and Hsp90ab1 (Fig2 C). Then, we knockdown Hdac6 in neurons to examine its role *in vivo* and the results showed that Hdac6 downregulation contributed to more neuronal death in tMCAO mice (Fig EV2B). Meanwhile, we also found that Hdac6-Kla level was elevated in cultured neurons under OGD/R (Fig 2D). To further verify the effect of Hdac6-Kla on neuron, we transfected Hdac6 plasmid with mutations in K915, k901, k896 and k888 (Hdac6-mutant) into primary cultural neurons to downregulate lactylation in Hdac6, and Hdac6-wildtype plasmid transfection was served as control. Hdac6 mutant plasmid effectively decreased the Hdac6-Kla level (Fig 2E). Next, we observed the effect of Hdac6-Kla in primary cultured neurons under OGD/R. The immunofluorescent and TUNEL results showed that inhibited Hdac6 lactylation induced evidently neuronal dendrites loss (Fig 2F) and a significant decrease in neuron viability (Fig 2G). These data indicated a protective role of Hdac6-Kla in cerebral ischemic reperfusion injury.

**Fig 2.**
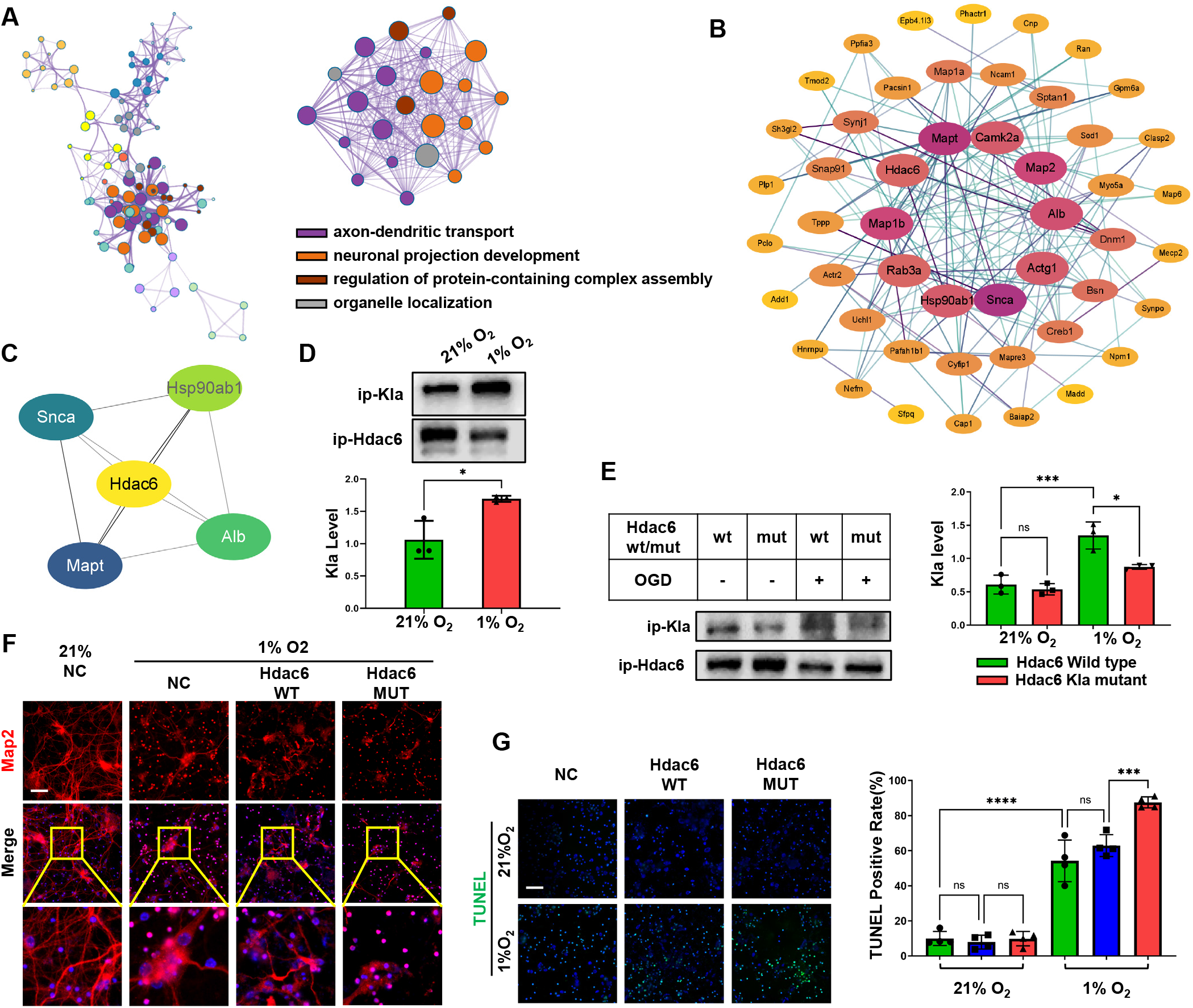
Global landscape of lactylatome and Functional analysis of lysine lactylation in ischemic stroke. A. The GO network of functional connections and clustering (Left). The top1 subset in quadratic clustering of GO network (Right). B. The protein-protein interaction (PPI) network of the top1 subset showed in panel A Right. C. The hub proteins identified from PPI network in panel B. D. Western blotting analysis of Hdac6 Kla in primary culture neurons after OGD/R. (n=3) E. Western blotting analysis of Hdac6 Kla in primary culture neurons with Hdac6 wild type or Hdac6 Kla-mutant plasmids after OGD/R. (n=3) F. Immunostaining of Map2 in primary culture neurons transfected with Hdac6 Kla-mutant plasmid after OGD/R (neurons with Hdac6 wild type plasmid as control). Scale bar: 100μm; G. TUNEL staining of primary culture neurons with Hdac6 wild type or Hdac6 Kla-mutant overexpression after OGD/R. Scale bar: 100μm; (n=4) Data in D are presented as mean ± SD (Student’s t-test); Data in E, G are presented as mean ± SD (Tukey’s multiple comparisons test); * P<0.05, *** P<0.001, **** P<0.0001, ns P>0.05.

### Hdac6-Kla modifies calcium homeostasis by binding immunoglobulin heavy chain binding protein

To further explore the potential downstream mechanisms mediated by Hdac6-Kla, proteomics analysis was conducted using Hdac6 immunoprecipitation protein sample from normoxia or hypoxia neurons. The heatmap in Figure EV3A showed that among the biological replicates within each group there were no significant differences in protein level. The volcano plot shows the 341 proteins identified (Fig EV3B). According to the cut-off value of 1.3 or 0.76 Foldchange, 85 down-regulated proteins and 97 up-regulated proteins were filtered (Fig. EV3C). The GO enrichment analysis showed the calcium signal pathway ranked within top10 (Fig3 A). Thereby, we conducted calcium staining in primary cultural neurons. The results indicated an intracellular calcium increase after OGD/R, which was aggravated by Hdac6 Kla mutant (Fig 3B). IP3 receptor inhibitor(2-APB), a specific endoplasmic reticulum (ER) calcium channel inhibitor, significantly abolished the increase of Hdac6 Kla on calcium (Fig 3B). Then we performed PPI network analysis for the whole identified proteins (Fig EV4A). Top 10 Hub proteins of the whole PPI network were Prkaca, Rab12, Mapk3, Hspa5, Mtor, Cdc42, Pten, Prkca, Grb2, Ywhag (Fig 3C). In order to clarify the correlation between protein function and calcium signal, we used the MCODE plugin(Bader & Hogue, 2003). to perform secondary clustering on the PPI network. Top 8 subsets were displayed (fig 3D left), the second-ranked subset of which is associated with calcium signaling downstream mechanisms (Fig 3D right). In this subset, Hspa5, encoding an immunoglobulin heavy chain-binding protein (BiP), was one of the top 10 hub proteins and was closely related to the regulation of calcium homeostasis in the endoplasmic reticulum. Then we transfected Hdac6-wildtype or Hdac6-Kla mutant plasmid by lentiviral vector to Neuro2a cell line after downregulating original Hdac6 by siRNA. Western blot analysis showed a reduction in Hdac6 Kla (Fig 3F) and an enhancement of Hdac6 binding BiP in Kla mutation (Fig 3G). Meanwhile, Hdac6 Kla mutant also affected the downstream endoplasmic reticulum calcium (ER) homeostasis-related proteins of BiP (Fig 3G). Downregulating BiP in Neuro2A eliminated the modulations of Hdac6 Kla on intracellular calcium accumulation (Fig 3E) and abrogated the modulations of endoplasmic reticulum calcium homeostasis regulatory proteins (Fig 3G). In summary, these data suggest that Hdac6 Kla regulated neuronal ER-associated calcium homeostasis, which may be related to the Hdac6-BiP interaction under hypoxic reoxygenation injury.

**Fig 3.**
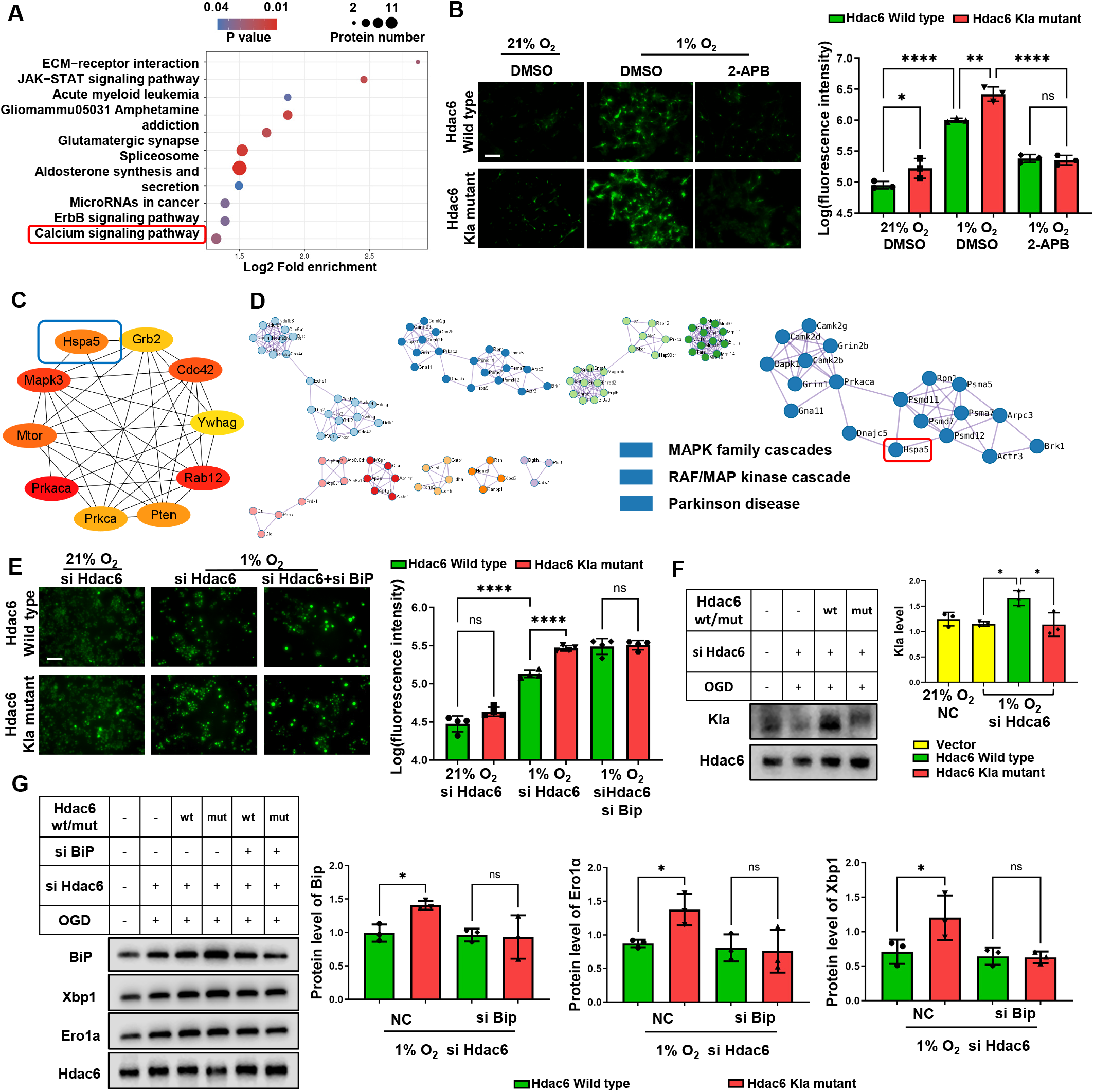
Hdac6-Kla mediates calcium overload by binding BiP. A. KEGG analysis of proteins binding Hdac6 in neurons after OGD/R. B. Intracellular calcium staining by Fluo-4 dye of primary culture neurons with Hdac6 wt/Kla-mutant after OGD/R. Scale bar: 100μm; (n=3) C. Hub proteins from the PPI network of Hdac6 binding proteins in neurons after OGD/R. D. Secondary clustered by MCODE plugin in the PPI network of Hdac6 binding proteins in neurons after OGD/R; E. Intracellular calcium staining by Fluo-4 dye in Neuro2A cell line co-transfected with BiP siRNA and Hdac6 Kla mutant or Hdac6 wild type plasmids after OGD/R. Scale bar: 100μm; (n=3) F. Western blotting analysis of Hdac6 Kla in in Neuro2A cell line co-transfected with BiP siRNA and Hdac6 Kla mutant or Hdac6 wild type plasmids after OGD/R; NC: Negative control; (n=3) G. Protein level of Hdac6 binging BiP, Ero1α and Xbp1 in Neuro2A cell line co-transfected with BiP siRNA and Hdac6 Kla mutant or Hdac6 wild type plasmids after OGD/R; NC: Negative control; (n=3) Data in B, E, F, G are presented as mean ± SD (Tukey’s multiple comparisons test); **\*** P<0.05, **\*\*** P<0.01, **\*\*\*** P<0.001, **\*\*\*\*** P<0.0001, ns P>0.05.

## Discussion

In current study, we found that patients with better neurological prognosis (mRS≤2) exhibited a higher serum lactate level. Then we demonstrated that inhibiting endogenous lactate increased infarct volume and aggravated neuronal damage accompanied with a decrease of protein lactylation in mice tMCAO model. The neuroprotection of endogenous lactate was also confirmed *in vitro*. Further, LC-MS/MS Analysis highlighted that Hdac6 lactylation increased in response to ischemic brain injury. Mutation of the lactylated lysine in Hdac6 exacerbated neuron damage under hypoxic condition. After performing immunoprecipitation proteomics, we found that reduction in Hdac6 lactylation enhanced its binding to BiP, an ER-localized protein that modulates calcium homeostasis, thereby leading to cytoplasmic calcium increase and ultimately exacerbating neuronal damage. Taken together, our present data suggested a new neuroprotection mechanism produced by endogenous lactate-derived Hdac6 lactylation via regulating calcium homeostasis, which may facilitate the development of new therapeutic strategies.

Lactate, a star in brain glucose metabolism, has been extensively studied as an energy resource for decades and risen controversies (Dienel, 2019). In cerebral ischemic stroke, the effects of lactate are also elusive and debatable. Some studies indicated that the lactate accumulation resulting from decrease cerebral blood would cause a toxic acidosis, while others found that lactate could be utilized as a fuel to support brain function (Annoni *et al*, 2021, Daniele *et al*, 2021). We found that patients with better prognosis have a higher serum lactate after vascular recanalization. Of note, the results still need a well-designed and large-sample clinical trial in further study, similar to the ongoing clinical trial, *Exploring the effect of lactate administration after ischemic stroke on brain metabolism (LacAVC)*, proposed by Professor Lorenz Hirt (MD, 2021). Apart from the original metabolic role, lactate has been newly found to enhance histone lactylation, an astonishing non-metabolic effect of lactate on post-translational modification, which is confirmed to regulate gene expression in macrophage (Zhang *et al*, 2019b), neuron (Hagihara *et al*, 2021), and microglia (Pan *et al*, 2022). In current experiments, we found that pan-lactylation is elevated after cerebral ischemic reperfusion and the lactylation is decreased when we inhibited the endogenous lactate, which is consistent with the previous finding by Zhang et al. showing that endogenous lactate is a crucial determinant of histone Kla levels. Interestingly, we also found that reduced lactylation due to reduced lactate led to larger infarct volumes and severer neuronal damage, suggesting that lactylation has a neuroprotective role in cerebral ischemia-reperfusion injury.

To further test our hypothesis, we performed LC-MS/MS analysis and detected lactylation variation in 204 proteins, the functional analysis of which suggested that the core functions were concentrated in regulations of axon-dendrites transportation and neuronal projection. These functions affected the morphology of neuronal axons and dendrites and were also critical for neuronal survival. We then found that lactylation of Hdac6 was upregulated in response to cerebral ischemia-reperfusion injury without significant changes in histone lactylation. Although previous studies have found that histone lactylation epigenetically regulated genes expression to affect the development of diseases, non-histone lactylation is critical in diseases by regulating the function of key proteins. HMGB1 lactylation resulted in more HMGB1 exocytosis and aggravating sepsis (Yang *et al*, 2022). Moreover, our present data demonstrated that Hdac6 lactylation regulates dendritic morphology and neuron survival after stroke. Previous studies have reported Hdac6 as one of the HDACs class IIb family members modulated neuronal damage in neurological diseases including ischemia stroke (Wang *et al*, 2016). One of the essential modifications of Hdac6 is deacetylation of cytoskeletal proteins α-tubulin, which destroys the stability of microtubules and consequently causes vesicle-transport dysfunction in neurons (Osseni *et al*, 2020). Another mechanism is the blockage of transport in mitochondria at neuron synapses, resulting in dysfunction of neuron synapses (d’Ydewalle *et al*, 2011). These key functions are similar to the functional alterations of lactylation HDAC6 in our study. Additionally, Hdac1-3 of the HDACs family have been found to work as delactylation enzymes (Moreno-Yruela *et al*, 2022). However, lactylation of HDACs family members themselves has not yet been reported. Our present data first demonstrated that Hdac6 can also be lactylated in lysine residues after stroke. We preliminarily demonstrated that the function of Hdac6 lactylation was associated with regulating ER-associated intracellular calcium homeostasis by BiP. Apart from above mentioned effect, Hdac6 can also modulate the oxidative stress level (Su *et al*, 2016) and inflammatory state of cells (Magupalli *et al*, 2020) by regulating protein deacetylation or ubiquitination (Balmik *et al*, 2020). Therefore, the functions of Hdac6 lactylation, including intervening cellular homeostasis, interacting with other PTMs, or modulating genes expression, remain to be explored in our following studies.

In conclusion, data from both stroke patients and mouse models implied the significance of lactate in neurological outcomes. Hdac6-Kla targeting BiP-related calcium homeostasis provides a new insight into lactate mechanism research, and sheds new light on the interpretation for the mechanism of injury in stroke. Hdac6-Kla is also expected to be a potential therapeutic target for neurological damage in ischemic stroke.

## Method and materials

### Clinical data acquisition

In the current study, we included 30 patients diagnosed with ischemic stroke, and 4 healthy volunteers as control in the First Affiliated Hospital of Jinan University. The ethical approval for the study was obtained by IRB of the First Affiliated Hospital of Jinan University (approval number: KY-2022-027). All stroke patients included underwent revascularization. And the venous blood was drawn from the patient immediately after revascularization, while venous blood samples of the healthy volunteers were taken the day signing the consent form. We assessed neurological outcome in stroke patients using the modified Rankin Scale (mRS) at 90 days, on a scale from 0 (asymptomatic) to 6 (death).

### Animals

Male C57/BL6 mice (22 ± 2g, Specific Pathogen Free, 8 weeks old) were obtained from the central animal facility of Southern Medical University (Guangzhou, China). The animals were housed under standard conditions of light and dark cycles (12 h:12 h, temperature 25 °C) with free access to food and water. And the cages were regularly cleaned. All experimental procedures were performed according to the approved protocols and guidelines of the Institutional Animal Ethical Care Committee of Southern Medical University Experimental Animal Centre.

### Transient middle cerebral artery occlusion model establishment

The transient middle cerebral artery occlusion (tMCAO) model was performed following a previously reported method in our article(Cao *et al*, 2021). Briefly, mice were anesthetized with continuous inhalation of sevoflurane (1.5%-3%). The model was established by inserting a monofilament from the external carotid artery to the middle cerebral artery. After the monofilament was inserted for 1.5 hours of ischemia, the monofilament was gently pulled out to form reperfusion. Mice subjected to tMCAO were randomly divided into each group. Oxamate (Selleck) was administered 500mg/kg intraperitoneally (Qu *et al*, 2021) at the onset of reperfusion. Mice in the sham group were injected with an equal amount of DMSO.

### Infarct Volume Analysis

2,3,5-Triphenyltetrazolium chloride staining (TTC (Sigma) was used to reflect cerebral infarction as a percentage of brain volume. The mice were anesthetized and the integral brains were quickly obtained and cut into 2 mm tissue slices, then stained with 2% TTC for 10 minutes and soaked in 4% formaldehyde for 24 hours. The brain slices were arranged in order and photographed. The area of cerebral infarction was calculated using the Image J 1.52a. The infarct area was calculated as the area of the non-ischemic hemisphere minus the non-infarcted area of the ischemic hemisphere. Infarct volume = infarct area × thickness (2 mm). The percent of cerebral infarction was calculated using the following formula: The percentage of cerebral infarction = infarct volume/the volume of the non-ischemic hemisphere×100%.

### Primary neuron culture

Cortical neurons were harvested from the cortices of C57Bl/6J mouse embryos (E16-18). Procedures and protocols have been approved by the Southern Medical University Administrative Panel on Laboratory Animal Care, and experiments were conducted in accordance with the guidelines of Animal Use and Care of Southern Medical University. After pregnant mice were anesthetized, embryos were removed from the uterus. Then embryonic cortices were dissociated mechanically into single cells. Cells were resuspended in Neurobasal Medium (Invitrogen) supplemented with B27 (Invitrogen) and Glutamax (Invitrogen). The cell suspension was plated on poly-L-ornithine and laminin-coated plates (PLL, Sigma-Aldrich, United States) and maintained in a 5% CO_2_ incubator at 37□. Half of the neuron media was replaced with fresh medium every three days.

### Neuro2a cell cultures

Neuro2a cells were cultured in DMEM (Gibco) supplemented with 10% fetal bovine serum (Gibco) in a 5% CO_2_ incubator at 37□. Cells were regularly passaged every two days.

### Mutation assay

Lactylation in K915, K901, K896 or K888 of Hdac6 were identified by lactylated LC/MS analysis. Accordingly, mutation of *hdac6* plasmid was conducted in genes encoding K915, K901, K896 and K888 leading to a lysine (K) to Arginine (R) mutation (Fig EV5A). The mutant sequence or wild-type sequence of HDAC6 were respectively cloned on the overexpression plasmid vector Lv206 (GeneCopoeia) (Fig EV5B). In order to package this vector into lentivirus, Lenti-Pac™ HIV Expression Packaging Kit (GeneCopoeia) was used for transfection in 293T cell line following the instructions of the kit. Pseudo virus-containing culture medium was collected in sterile capped tubes at 48 hours and 72hour post transfection respectively and centrifuged at 500 x g for 10 minutes to get rid of cell debris. Following centrifugation, the supernatant was filtered through 0.45 µm polyethersulfone (PES) low protein-binding filters.

### Cell transfection

Cells were transfected siRNA using lipofectamine 3000 (Invitrogen) reagents following the manufacture’s guidelines. After adding the transfection reagent for 6h, the mediums were discarded and replaced by fresh mediums. The sequences of siRNA for Hdac6 and BiP were shown in Table 1. Further experiments were conducted after 48 h after transfections. Lentivirus with Hdac6 mutant or Hdac6 wildtype plasmid were applied to neuron and Neuro2a cell line for 24hours. Virus-containing medium was replaced with fresh medium. Further experiments were conducted after 48 h after transfections.

### Cell treatment

Cells were subjected to oxygen and glucose deprivation followed by reoxygenation (OGD/R), to mimic the ischemic/reperfusion-like condition in vitro. For neurons, the original media was collected, saved and replaced with glucose-free DMEM. Then the neurons were transferred to a hypoxic incubator continuously gassed with 95% N_2_, 5% CO_2_ and 1% O_2_ at 37□for 4 hours. Then the glucose-free medium was replaced with the pre-collected Neurobasal Medium and the cells were then returned to a normoxic incubator. The control cells were kept in Neurobasal Medium in normoxic incubator. To model OGD/R in N2a cells, the media were replaced with glucose-free DMEM and saturated with 5% CO_2_/95% N_2_ for 12 hours. Oxamate was dissolved in doble distilled water and diluted with culture medium to 20mM. The cells were subjected to the drug for 6 hours and the medium was replaced with fresh medium. The cells were harvested at 6 hours after reoxygenation.

### TUNEL assay

Neuron apoptosis was detected by TUNEL staining according to the instructions of the kit (Vazyme). Cells were fixed and permeabilized with 0.2% Triton X-100. Subsequently, cells were added with labeling mixture containing FITC-12-dUTP at 37□for 1h. After washed for 3 times, cells were mounted with the DAPI mounting medium (Solarbio).

### Immunoprecipitation

The cells were lysed with RIPA lysis buffer containing protease inhibitors on ice for 30 minutes. After centrifugation (12,000 g for 20 min at 4□), the supernatant was mixed with anti-Hdac6 antibody(Abcam, 1:100)overnight at 4□to form immunocomplexes. Then 1/10 volume of protein A+G magnetic beads (Beyotime) were added to the immunocomplexes and incubated for 6 hours at 4□with mixing. Afterwards, beads were collected with a magnetic stand and washed three times with RIPA lysis buffer. 60µL of sample loading buffer was added to the beads. Then the mixture was heated at 96-100ºC in a heating block for 8 minutes. Magnetically separate the beads and save the supernatant containing the target antigen for western blotting analysis.

### Immunofluorescence staining

At 24h after tMCAO, mice were deeply anesthetized and perfused intracardially with saline and 4% paraformaldehyde (PFA, Solarbio). After fixation in 4% PFA, the brains were dehydrated in 30% sucrose and cut into frozen coronal slices (40µm). Brain slices were permeabilized with 0.3% TritonX-100 (Sigma-Aldrich) and blocked by 5% bovine serum albumin BSA (Solarbio) for 60 min at room temperature. Then the slices were incubated overnight at 4□with the primary antibodies anti-L-lactyl lysine (1:250 dilution; rabbit, PTM Bio) and anti-Map2 (1:250 dilution; rabbit, Proteintech). Thereafter, the brain slices were washed with PBS and were incubated with the appropriate fluorescent labelled secondary antibodies for 1 hour at room temperature. After washing, brain slices were counterstained with DAPI (Solarbio, China) and analyzed using laser-scanning confocal microscopy (LSM900, Japan).

Primary cultural neurons were washed once with PBS, fixed for 30 minutes in 4% paraformaldehyde (Solarbio, China) at 37 □and permeabilized with 0.5% Triton X-100 (Sigma-Aldrich) for 10 minutes. After 5 min-wash with PBS three times, the cells were blocked with 3% BSA (Solarbio, China) for 1 hour at room temperature. Cells were then incubated overnight at 4 □with primary antibodies against map2 (1:300 dilution; Proteintech; China). After washed the cells were incubated for 1 hour at room temperature with fluorescent labelled secondary antibodies. After washing, the cells were counterstained with DAPI (Solarbio, China) and analyzed using laser-scanning confocal microscopy (LSM900, Japan).

### Lactate Colorimetric Assay

The serum lactate from cerebral ischemia stroke patients and healthy volunteers was measured by the Lactate Assay Kit (Sigma) according to the manufacturer’s protocol. Optical density was detected at 530nm wavelength using Biored microplate reader.

### LC-MS/MS Protein Sample Preparation

The brain sample was grinded with liquid nitrogen into cell powder. Lysis buffer (8 M urea, 1% protease inhibitor cocktail) was then added to the cell powder, followed by sonication three times on ice using a high intensity ultrasonic processor (Scientz). The remaining debris was removed by centrifugation at 12,000g at 4 □for 10 min. Finally, the supernatant was collected and the protein concentration was determined with BCA kit according to the manufacturer’s instructions. The protein solution was reduced with 5mM dithiothreitol for 30 min at 56 □and alkylated with 11mM iodoacetamide for 15min at room temperature in darkness. The brain sample was then diluted by adding 100 mM TEAB to urea concentration less than 2M. Finally, trypsin was added at 1:50 trypsin-to-protein mass ratio for the first digestion overnight and 1:100 trypsin-to-protein mass ratio for a second 4h-digestion. Finally, the peptides were desalted by C18 SPE column. Pan-antibody-based PTM enrichment: to enrich modified peptides, tryptic peptides dissolved in NETN buffer (100 mM NaCl, 1 mM EDTA, 50 mM Tris-HCl, 0.5% NP-40, pH 8.0) were incubated with pre-washed antibody beads (anti-L-lactyl lysine, PTM Bio) at 4□overnight with gentle shaking. Then the beads were washed for four times with NETN buffer and twice with DDH_2_O. The bound peptides were eluted from the beads with 0.1% trifluoroacetic acid. Finally, the eluted peptides were combined and vacuum-dried. The resulting peptides were desalted with C18 ZipTips (Millipore) for LC-MS/MS analysis.

### 4D Mass Spectrometer

The tryptic peptides were dissolved in solvent A (0.1% formic acid, 2% acetonitrile/in water), and separated with a gradient from 6% to 24% solvent B (0.1% formic acid in acetonitrile) over 70 min, 24% to 35% in 14 min and climbing to 80% in 3 min then holding at 80% for the last 3 min, all at a constant flow rate of 450 nL/min on a nanoElute UHPLC system (Bruker Daltonics). The peptides were subjected to capillary source followed by the timsTOF Pro mass spectrometry (Bruker Daltonics). The electrospray voltage applied was 1.60 kV. Precursors and fragments were analyzed at the TOF detector, with a MS/MS scan range from 100 to 1700 m/z. The timsTOF Pro was operated in parallel accumulation serial fragmentation (PASEF) mode. Precursors with charge states 0 to 5 were selected for fragmentation, and 10 PASEF-MS/MS scans were acquired per cycle. The dynamic exclusion was set to 30s. The identified proteins of lactylation and Hdac6 immunoprecipitation were list in table1 and table2.

### Database Search

The resulting MS/MS data were processed using MaxQuant search engine (v.1.6.15.0). Tandem mass spectra were searched against the human SwissProt database (20422 entries) concatenated with reverse decoy database. Trypsin/P was specified as cleavage enzyme allowing up to 2 missing cleavages. The mass tolerance for precursor ions was set as 20 ppm in first search and 5 ppm in main search, and the mass tolerance for fragment ions was set as 0.02 Da. Carbamidomethyl on Cys was specified as fixed modification, and acetylation on protein N-terminal and oxidation on Met were specified as variable modifications. FDR was adjusted to < 1%.

### Functional enrichment analysis

Functional enrichment analysis and the functional networks of the identified proteins was constructed by Metascape (https://metascape.org (Zhou *et al*, 2019). To find out the densely connected regions that were closely related to cerebral ischemia-reperfusion injury, secondary clustering of the functional network was conducted by the molecular complex detection (MCODE), a plug-in module of Cytoscape (Bader & Hogue, 2003). The PPI analysis was performed through STRING version 11.5 (https://string-db.org/). The hub proteins from the PPI network were predicted by the cytoHubba, one of the plugins of Cytoscape (Chin *et al*, 2014).

### Statistical analysis

Statistical analysis was conducted using Graphpad Prism 9. Differences were evaluated by t test (nonparametric t test) or one-way (Bonferroni’s multiple comparisons test). p < 0.05 was considered statistically significant.

## Data availability

The data that support the findings of this study are available from the corresponding author, upon reasonable request.

## Acknowledgments

This work was funded by the National Natural Science Foundation of China (Grant Number: 81973305), Natural Science Foundation of Guangdong Province, China (Grant Number: 2021A1515010897), the Discipline Construction Fund of Central People’s Hospital of Zhanjiang (Grant Number: 2020A01 and 2020A02) and supported by Shenzhen Fundamental Research Program, China (Grant Number: JCYJ20190809171003698).

## Author contributions

**Jun Cao:** Conceptualization; Formal Analysis; Investigation; Methodology; Writing original draft. **Zexuan Hong:** Conceptualization; Formal Analysis; Investigation; Methodology; Writing original draft. **Fanning Zeng:** Investigation. **Zaisheng Qin:** Investigation. **Miao Lin:** Clinical data acquisition. **Hao Wang:** Clinical data acquisition. **Daming Zuo:** Supervision; Funding acquisition; Writing-review& editing. **Tao Tao:** Supervision; Funding acquisition; Writing-review& editing.

## Disclosure and competing interests statement

The authors declare that they have no conflict of interest.

## Notes

### Competing Interest Statement

The authors have declared no competing interest.

